# Thermostability improvement of *Aspergillus awamori* glucoamylase via directed evolution of its gene located on episomal expression vector in *Pichia pastoris*

**DOI:** 10.1101/652164

**Authors:** Alexander Schmidt, Alexey Shvetsov, Elena Soboleva, Yury Kil, Vladimir Sergeev, Marina Surzhik

## Abstract

Novel thermostable forms of glucoamylase (GA) from filamentous fungus *Aspergillus awamori* X100 were constructed using the directed evolution approach based on random mutagenesis by error-prone PCR of the catalytic domain region of glucoamylase gene with its localization on a new episomal expression vector pPEHα in *Pichia pastoris* cells. Of 3000 yeast transformants screened, six new thermostable GA mutants with amino acid substitutions Val301Asp, Thr390Ala, Thr390Ala/Ser436Pro, Leu7Met/His391Tyr, Asp9His/Ile82Phe, Ser8Arg/Gln338Leu were identified and studied. To estimate the effect of every single substitution in the double mutants, we have constructed an appropriate single mutants of GA by site-directed mutagenesis and analyzed their thermal properties. Results of the analysis showed that only Ile82Phe and Ser8Arg mutations caused an increased thermostability. While Leu7Met and Asp9His mutations decreased the thermal stability of GA, and Gln338Leu had little effect, the synergistic effect of double mutant forms Leu7Met/His391Tyr, Asp9His/Ile82Phe and Ser8Arg/Gln338Leu revealed the significant thermostability improvement as compared to wild type GA.

## Introduction

Glucoamylase (EC 3.2.1.3, 1,4-α-D-Glucan glucohydrolase) is an important enzyme applying for starch saccharification in the industrial glucose and bioethanol production. The enzyme cleaves 1,4-α-glycosidic bonds from the nonreducing end of the glycosidic chains of starch, glycogen and maltooligosaccharide releasing free β-D-glucose. Glucoamylase (GA) from filamentous fungus *Aspergillus awamori* X100 possesses an optimal activity at 55-60°C with the irreversible thermal inactivation at higher temperature. In biotechnological processes of raw starch conversion two other much more thermostable enzymes, α-amylase for primary hydrolysis and liquefying of starch and glucoisomerase to convert some of glucose into fructose are used. These three enzymes are commonly applicable for the production of glucose-fructose syrup, which is widely used in food and bakery industry. The obtaining of GA forms with improved characteristics, including increased thermal stability of enzyme, would be commercially useful for applying in the optimized technological processes.

Starch degrading enzymes including GAs are widely distributed among different microorganisms. The most active forms were found in a number of moulds and yeast, but new source of GA with potentially applicable properties is still of a considerable attention [1]. However, the industrial focus has been on GA from *Aspergillus* genus and *Rhizopus oryzae* [2].

To increase the thermostability of GA different approaches of protein engineering, including site-directed mutagenesis based on computer modeling [3–8] and molecular dynamics [9, 10] have been described. These studies were stimulated by solving the crystal structure of GA from *Aspergillus awamori* X100 in the early 90^th^ [11, 12]. But the most successful results were achieved in the works, where the method of directed evolution was used [13, 14].

In present work, we reported the novel thermostable forms of GA from *A. awamori* X100, obtained by directed evolution of its gene with episomal localization in *Pichia pastoris* cells, and analyzed their activities and thermostability. Besides we discussed the cases of synergistic combination of single amino acid substitutions and proposed an explanation on the significant thermostability improvement for these double mutants of GA.

## Materials and Methods

### Plasmids, Strains and Media

Episomal expression vector pPEHα have been constructed to make easy the procedure of putative mutant selection in *Pichia pastoris;* details, plasmid inheritance stability and rate of GA expression will be described separately. Briefly, plasmid contains origin of replication *PARS1* from *Pichia pastoris, HIS4* gene form *S. cerevisiae* for histidine selection, *AOX1* promoter and alpha-factor signal peptide. Ampicilin at concentration of 100 mkg/ml in LB solid or liquid medium was used to select clones and harvest *E. coli* DH5α cells for DNA manipulations. *P. pastoris* GS115 cells were grown on standard YPD medium while transformants bearing pPEHα plasmid or its variants were grown on Pichia minimal growth medium (MM, MDS or MG). Plasmid DNA and DNA libraries were transformed in DH5α or GS115 by electroporation [15–17].

### Cloning of *A. awamori* X100 glucoamylase gene into pPEHα

Full length gene of *A. awamori* X100 glucoamylase GlaA was amplified by PCR with Pfu DNA polymerase using YEpPM18 plasmid [18] as a template. The following oligonucleotides: GlaA-fw (5’-aaactcgagaaaagagcgaccttggattcatgg-3’) and GlaA-rv (5’-acatctagacgaaatggattgattgtc-3’) were used. The PCR product was digested by XhoI and XbaI endonuclease and cloned into pPEHα vector with the same restriction sites. The resulting plasmid pPEHα-GlaA (Fig. 1) was transformed in *P. pastoris* GS115 with the selection of transformants by their ability to grow on histidine-deficient medium.

**Figure 1.**
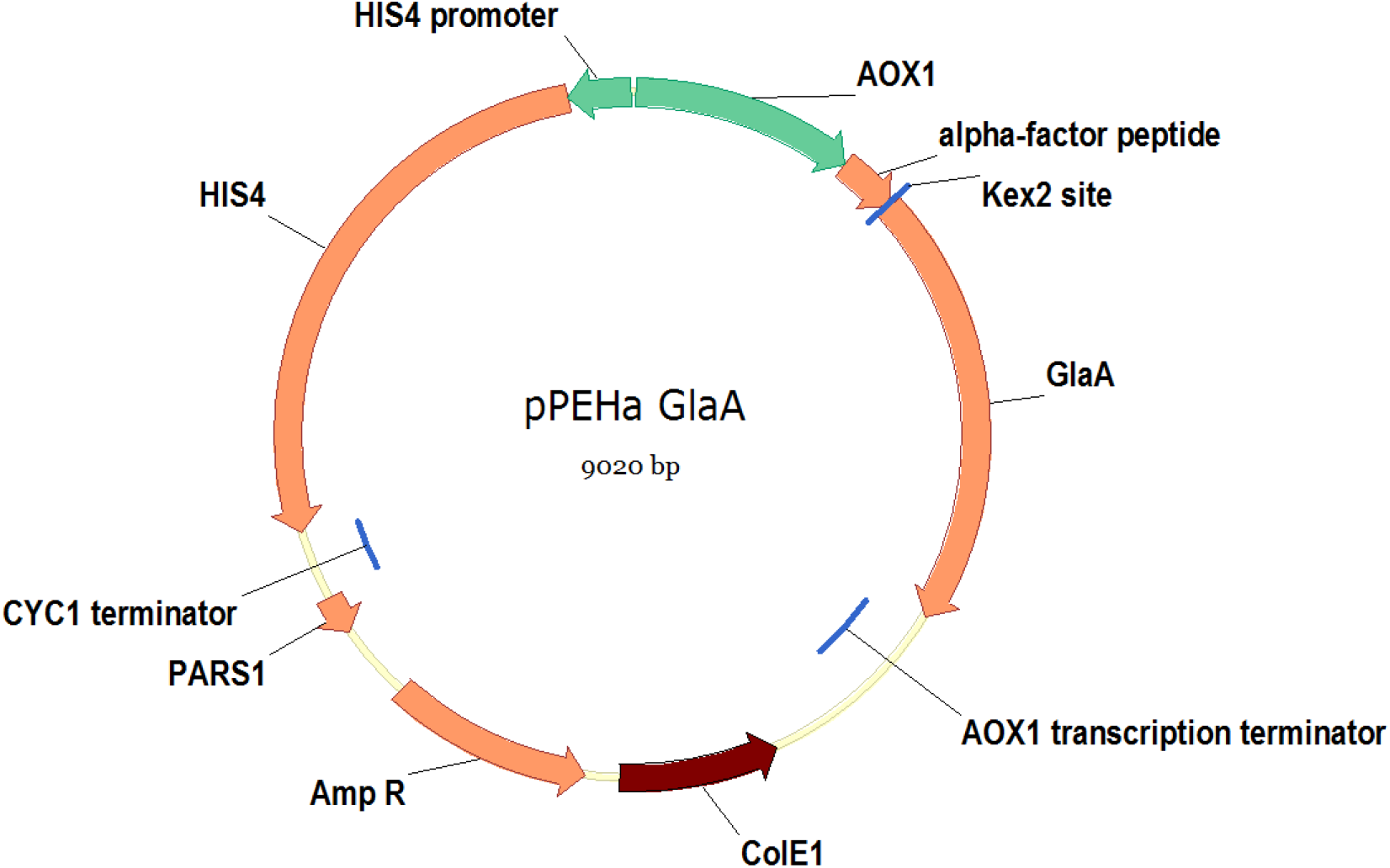
Scheme of the episomal expression plasmid pPEHα-GlaA.

### Site-directed mutagenesis

Mutagenesis was performed by method of partially overlapped primers [19, 20] using iProof DNA-polymerase (Bio-Rad). The PCR reaction mixture was digested with DpnI, precipitated by ethanol and transformed in *E. coli* DH5α cells by electroporation. Mutations were confirmed by allele-specific PCR [15] and Sanger sequencing.

### Random mutagenesis and Library construction

Random mutagenesis of GA gene fragment corresponding to the catalytic domain of glucoamylase (GA-CD) was performed by error-prone PCR based on increased concentrations of dCTP and dTTP nucleotides in reaction, and in the presence of magnesium chloride [21]. Each 50 μl of reaction mixture contained 50 ng of pPEHα-GlaA plasmid DNA as template, 5 unit of Taq-polymerase, 0.3 μM of each oligonucleotide GlaA-fw (5’ aaactcgagaaaagagcgaccttggattcatgg 3’) and GAkat-rv (5’ gtcttgctggtcgaggtcacgctgc 3’), 7 mM MgCl_2_, 0.5 mM MnCl_2_, 10 mM Tris-HCl (pH 9.0), 50 mM KCl, 0.1% Triton X-100, and deoxynucleoside triphosphates at the following concentrations: 0.2 mM dGTP, 0.2 mM dATP, 1 mM dCTP, 1 mM TTP.

After digestion by XhoI and BamHI endonucleases the mutated PCR-fragments was purified and ligated into linearized by the same restriction sites pPEHα-GlaA plasmid. The ligase reaction was transformed in *E. coli* DH5α cells to produce a library of mutagenized GA genes.

### Primary screening of transformants with the active GA forms

To identify the transformants with active GA, they were grown on plates with MM medium containing 1% methanol and 0.05% soluble starch at 30°C for 3 days. The yeast colonies with secreted GA activity formed “halos” around them in plate medium revealed by staining with iodine vapour.

### Selection of clones expressing improved glucoamylase

The selection of clones that expressed mutant variants of glucoamylase with increased thermostability included several stages: expression of the enzyme in a 96-well plate, determination of glucoamylase activity in the culture fluid, thermal inactivation, determination of residual activity, determination of thermal stability.

#### Expression

The glucoamylase activity positive clones were transferred into a 96-well culture plate with 180 μl of MM expression medium in each well. As a control, wild-type glucoamylase producing clone was transferred to each plate. The cells were incubated in a shaker-thermostat at 30°C, 250 rpm for 72 hours. Then the plates were centrifuged at 1100 g for 10 minutes.

#### Activity assay

To determine the glucoamylase activity, 30 μl of the supernatant was taken from each well and transferred to a new 96-well titration plate. To each sample 30 μl of maltose at concentration of 10 mg/ml in 0.2 M sodium acetate buffer pH 4.5 was added, and the plates was incubated at 37°C for 1 hour. Then, 340 μl of the glucose oxidase mixture was added to each well to determine the glucose accumulated and incubated at 37°C for 20 minutes. The absorption A_act_ was measured on a plate photometer at a wavelength of 495 nm.

#### Thermal inactivation and determination of residual activity

Thermal inactivation was performed in an amplifier in 96-well PCR plates. The procedure was as follows: 40 μl of the supernatant was taken from each well of the culture plate and added to 4 μl of 1 M sodium acetate buffer pH 4.5. The enzyme was inactivated at 75°C for 5 minutes with further incubation at 4°C for 20 minutes.

To determine the residual activity, 30 μl of each inactivated sample was transferred to a new titration plate and mixed with 30 μl of starch with a concentration of 10 mg/ml in 0.1 M sodium acetate buffer (pH 4.5). The plate was incubated at 37°C for 20 hours, and then 340 μl of the glucose oxidase mixture was added to each well and incubated at 37°C for 20 minutes. The absorption value of A_inact_ was measured on a plate photometer at a wavelength of 495 nm.

#### Thermal stability determination

For each sample, the ratio A_inact_/A_act_ was calculated. The mutant variant of glucoamylase was considered thermostable if the A_inact_/A_act_ ratio was greater than that of the wild-type enzyme.

Transformants producing the putative glucoamylase mutant forms with enhanced thermostability were re-expressed in 2 ml of culture medium in a 24-well plates at 30°C, 250 rpm for 72 hours. Then the culture fluid was subjected to inactivation at 75°C for 5, 6 and 7 minutes, after that the results were compared with the data of the primary selection.

### Glucoamylase purification

The yeast transformants was incubated in a shaker-thermostat in a volume of 8-16 ml of liquid yeast minimal medium at a temperature of 30°C, 250 rpm for three days. Methanol was added to 1% final concentration every 24 hours.

At the end of the cultivation, the medium was cooled to 6°C and centrifuged at 6000 rpm for 20 minutes. The supernatant was dialyzed against 50 mM Tris-HCl buffer pH 7.5 with 50 mM NaCl. Ion exchange chromatography was performed on a MonoQ 5×50 mm column (Pharmacia). Glucoamylase was eluted with a salt gradient from 50 to 500 mM NaCl in 50 mM Tris-HCl pH 7.5. Glucoamylase was identified by glucose accumulation in samples using a standard glucose oxidase system (ECO-service, Russia). Containing glucoamylase fractions were combined and dialyzed against 50mM NaOAc buffer pH 4.5 containing 50 mM NaCl.

### Enzymatic activity determination

To determine the enzymatic activity, the amount of glucose accumulated during enzymatic hydrolysis of maltose in 0.1 M sodium acetate buffer at pH 4.5 and temperature 50°C per unit of time at substrate concentrations >> K_M_ (40 mg/ml) was measured. From the reaction mixture of the enzyme and the substrate, samples of 10 μl were taken at certain intervals, cooled in liquid nitrogen, and analyzed for glucose using hexokinase or glucose oxidase systems (ECO-service, Russia). The increment of glucose concentration in the reaction mixture per unit of time characterized the activity of glucoamylase.

### Thermostability determination

The temperature dependences of the thermoinactivation constants were investigated as the main criterion for the thermal stability of the enzyme forms. The enzyme solution was incubated at a given temperature and samples were taken at certain intervals, which were immediately cooled in an ice bath and stored at 4°C for 24 hours. Then, in each sample the residual enzyme activity was determined at standard temperature of 50°C. The enzyme thermal inactivation constant, k_d_, was calculated as a slope of the straight line on the graph of 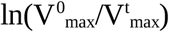 versus time. The amount of enzyme required to obtain 1 μm of glucose per minute was taken as unit of activity.

### Molecular Modeling

Glycam web server services, the PyMOL software package [22], as well as the GROMACS 2018 software package [23] were used for computer simulation. The structure of GA catalytic domain (PDB ID 1GAI) was used for the construction of the initial model. Using homologous modeling, the amino acid sequence errors in crystal structure were corrected. Using the GLYCAM server, weakly branched oligosaccharides each consisting of nine mannose residues and two N-acetylglucosamines were added at two N-glycosylation sites of glucoamylase.

### Molecular dynamics and ΔΔG^≠^ calculation

Molecular dynamics (MD) modeling was performed in the software package GROMACS 2018, while the amber99sb-ildn field [24] was used for the peptide part of the protein, glycam06h force field was used for glycosides [25] and tip3p was used as a water model. To calculate the change in ΔΔG^≠^ free energy of thermoinactivation caused by amino acid substitutions using the FEP method, the hybrid amino acid topology using PMX was built [26–28]. In the position of interest, the amino acid corresponding to the wild type GA was replaced by an amino acid with a hybrid topology, which contained the topologies A (corresponding to the wild type GA) and B (corresponding to the GA mutant form).

The systems were equilibrated separately for each state A and B. The resulting systems were placed in a periodic water box in such a way that at least 25Å remained to the walls of the box. Then the resulting box was minimized using the steepest descent algorithm. The system obtained as a result of minimization was subjected to charge neutralization by adding 50 mM of Na^+^ and Cl^−^ ions, while the total charge of the system was zero. The neutralized system was again subjected to the procedure of minimizing energy using the steepest descent algorithm. Next, the system was equilibrated using two stage approach. At the first stage, all heavy atoms of proteins and sugars were restrained to their initial positions using an additional energy term (posres), while at the start of equilibration the temperature (particle velocity distribution) was taken from the Maxwell distribution for a given temperature (348K was used). The system was equilibrated for 5 ns for each temperature. The integration step was 2 fs. At this stage, the Berendsen thermostat and the Berendsen barostat were used. At the second stage, the additional restraining potential was removed, and all components of the system could move freely. During this stage, the Nose-Hoover thermostat and the Parinello-Raman barostat were used, and the system was equilibrated for 10ns. Further, out of the last 5 ns of the second stage of equilibration, 101 states were selected in increments of 50 ps. For each state, the method of calculating the free energy based on FEP FastGrows algorithm was used to calculate the transition A → B and B → A for 100 ps with a step of 2 fs. The resulting 200 trajectories were analyzed using the CROOKS method described in [27] and implemented in PMX [28].

## Results

A 1.5-kb fragment corresponding to the catalytic domain of the enzyme (1-440 aa) was subjected to random mutagenesis by error-prone PCR and cloned again into pPEHα-GlaA to create the library of full-length GA with mutagenized GA-CD. Constructed in this manner DNA library of mutagenized forms of GA in the episomal expession vector pPEHα was screened by three-stage selection procedure for significantly increased thermal stability.

At first step, the DNA library was transformed in *P. pastoris* GS115 cells and colonies obtained were screened for ability to produce an active GA on plates with expression medium MM, containing 1% methanol and 0.05% starch. Among 3000 transformants analyzed, we selected 1000 colonies possessed the significant GA activity. In parallel all transformants were harvested on MG medium to maintain selected clones.

At second step, the 1000 selected mutants were incubated in 96-well plate with the liquid expression medium at 30°C in shaker-thermostat (New Brunshwick Scientific) at 250 rpm for 3 days. Primary selection of mutants with enhanced thermostability was as described in Materials and Methods followed by next round of the revealing the putative thermostable clones in 2 ml of medium in a 24-well plate for another 3 days. The amount of culture fluid obtained at this third step was sufficient to accurate determination of residual activity after incubation at 75°C for 5, 6 and 7 minutes. As a result, we found out 6 clones, secreted mutant variants of GA with considerably enhanced thermostability as compared with wild type GA. Sanger sequencing of plasmid DNA showed that there were two proteins with single substitutions Val301Asp and Thr390Ala, and four mutant forms with double mutations Thr390Ala/Ser436Pro, Leu7Met/His391Tyr, Asn9His/Ile82Phe and Ser8Arg/Gln338Leu. Notably, that only Ser436Pro and His391Tyr have been studied previously [29, 13]. It should be also noted that amino acid substitutions of Ser8Arg, Asn9His and Leu7Met occurred in α-helix 1, which seems to be one of the most thermosensitive regions in the protein structure [4, 8–10].

Table 1 shows the relative activities of GA mutant forms measured at pH 4.5 and 50°C using maltose as a substrate. As we can see, the Asn9His/Ile82Phe and Thr390Ala had increased activity, whereas the relative activity of Val301Asp, Thr390Ala/Ser436Pro and Ser8Arg/Gln338Leu were similar to wild-type GA. Variant Leu7Met/His391Tyr possessed a slightly reduced activity.

**Table 1.** Relative activity observed at pH 4.5 and 50°C using maltose as a substrate.

| GA | Relative activity |
| --- | --- |
| WT | 1 |
| Val301Asp | 1,16 |
| Thr390Ala | 1,38 |
| Thr390Ala/<br>Ser436Pro | 1,02 |
| Leu7Met/His391Tyr | 0,81 |
| Ile82Phe | 1,45 |
| Asp9His/Ile82Phe | 1,73 |
| Ser8Arg | 0,97 |
| Ser8Arg/Gln338Leu | 1,19 |

The thermal inactivation of wild-type and mutant forms of enzyme was studied at pH 4.5 and at seven temperatures in range from 65°C to 80°C. Semilogarithmic plotting of residual activity vs. inactivation time was used to determine the inactivation rate constant in accordance with first-order kinetics. Effects of temperature on thermoinactivation rate constant *k*_d_ for WT and mutant enzymes were presented on Fig. 2. All mutant variants appeared to be significantly more thermostable than wild-type GA and the difference became higher with the temperature increasing. Calculated values of the ΔΔG^≠^ are shown in Table. 2.

**Figure 2.**
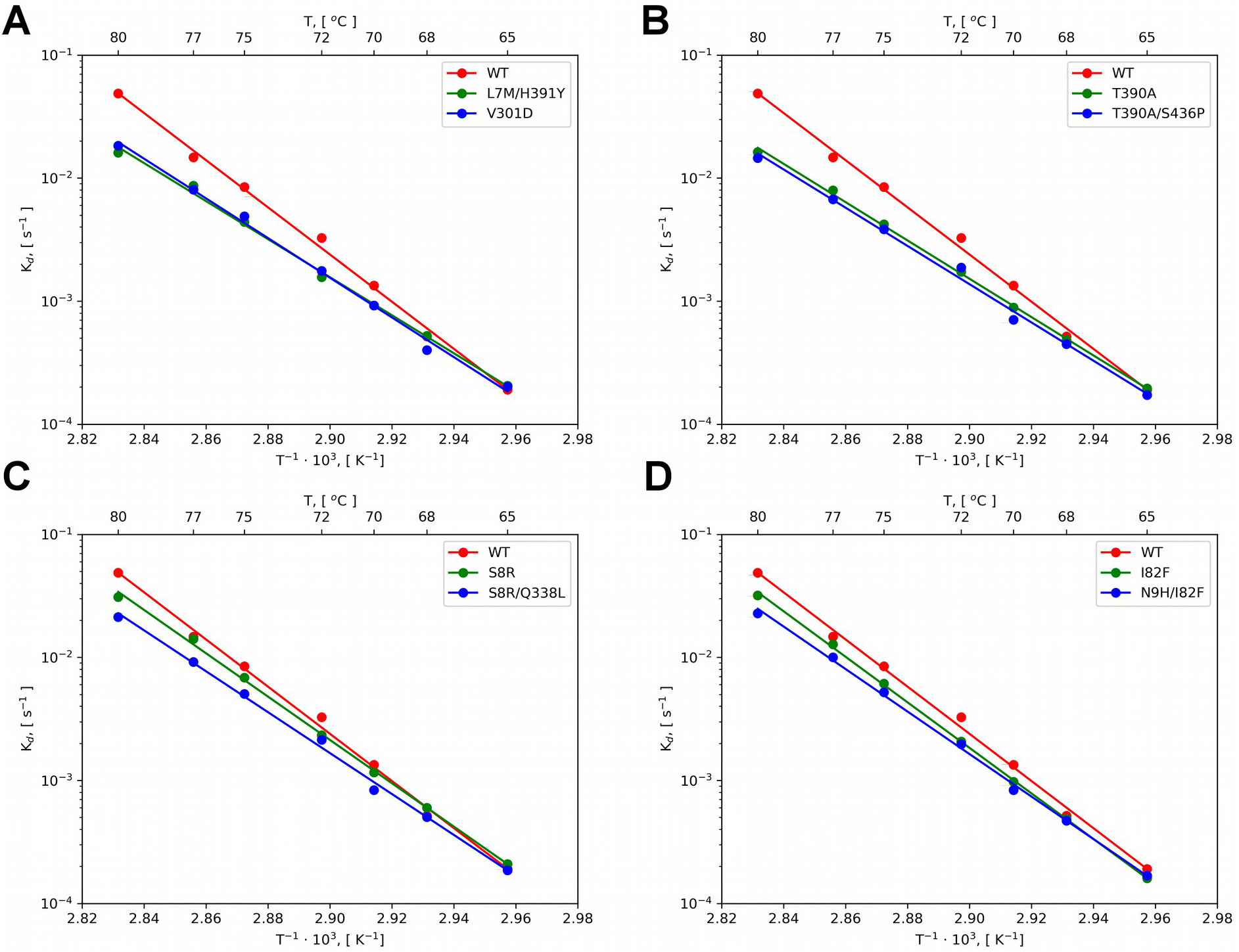
Effect of temperature on first-order thermal inactivation rate constants (k_d_): (A) Wildtype GA (WT) and mutant variants Val301Asp and Leu7Met/His391Tyr (B) GA WT and mutant variants Thr390Ala and Thr390Ala/Ser436Pro (C) GA WT and mutant variants Ser8Arg and Ser8Arg/Gln338Leu (D) GA WT and mutant variants Ile82Phe and Asn9His/Ile82Phe.

**Table 2.** Change of free activation energy of thermoinactivation (ΔΔG^≠^) of wild type and mutant variants of glucoamylase

| GA | $\Delta\Delta G^\ddagger$ (kJ/mol) | | | | Method / Reference |
| --- | --- | --- | --- | --- | --- |
|  | 70°C | 75°C | 80°C | CROOCKS<br>(75°C) |  |
| Val301Asp | 0.81 | 1.77 | 2.72 |  | DE |
| Thr390Ala | 0.96 | 1.98 | 2.99 | 6.80 | DE |
| Thr390Ala/<br>Ser436Pro | 1.22 | 2.16 | 3.1 |  | DE |
| Leu7Met | -1.83 | -2.15 | -2.44 | -2.48 | SDM |
| His391Tyr | - | - | 0.2<sup>&</sup> | -5.48 | <sup>&</sup> [13] |
| Leu7Met/His391Tyr | 0.79 | 1.78 | 2.76 | 5.61 | DE |
| Asp9His | -0.94 | -0.97 | -1.01 | -1.07 | SDM |
| Ile82Phe | 0.69 | 0.88 | 1.08 | 8.82 | SDM |
| Asp9His/Ile82Phe | 0.9 | 1.45 | 2.0 | 2.28 | DE |
| Ser8Arg | 0.18 | 0.62 | 1.06 | 3.71 | SDM |
| Gln338Leu | -0.52 | -0.53 | -0.54 | -0.80 | SDM |
| Ser8Arg/Gln338Leu | 0.79 | 1.51 | 2.23 | 5.73 | DE |

\*Activation energies of unfolding were calculated $\Delta G^\ddagger = -RT \ln(k_d h / K_B T)$ , where R is universal gas constant, h is Plank constant, $K_B$ is Boltzmann's constant, T is absolute temperature and $k_d$ – irreversible thermoinactivation constant [30]. DE – Directed evolution; SDM – Site directed mutagenesis.

In double mutants of GA, each amino acid substitution can add positive or negative effect on protein thermostability. To elucidate this influence we construct the Leu7Met, Asn9His, Ile82Phe, Ser8Arg and Gln338Leu single mutations in GA by site-directed mutagenesis. Results of the analysis showed that only Ile82Phe and Ser8Arg mutations caused an increased thermostability. While Leu7Met and Asp9His mutations decreased the thermal stability of GA and Gln338Leu had little effect.

Further study of the effect of these amino acid substitutions on the structure of GA was carried out by molecular dynamics methods, while the compliance of the calculated change of free energy ΔΔG^≠^ with the experimental data was used as another criterion for the correctness of the simulation results

Substitution Val301Asp led to a restructuring of the network of hydrogen bonds between amino acids 292, 301 and 302, and formation of new hydrogen bond between Asp301 side chain and Ala302 amine group, which led to stabilization of loop consisting of 292-302 amino acids (Fig. 3).

**Figure 3.**
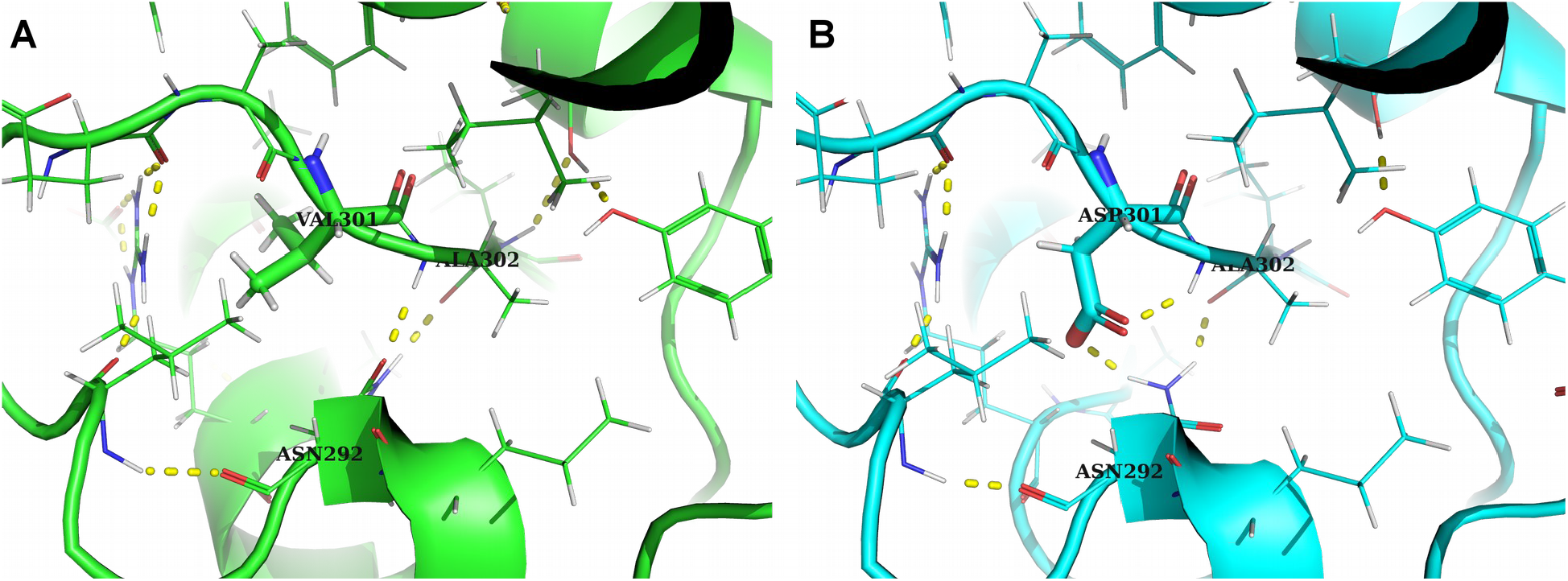
Amino acid substitution Val301Asp. The structures of the GA WT (A) and Val301Asp mutant variant (B) are indicated. The Val301Asp substitution led to a restructuring of the hydrogen bond network between amino acids 292, 301 and 302, and formation of a new hydrogen bond between Asp301 side chain and Ala302 amine group.

One of the most thermostable variants was the mutant with another single amino acid substitution Thr390Ala. Thr390 is located at the C-terminus of α-helix 12 (Fig. 4A), followed by an extensive loop containing the catalytic amino acid Glu400. Analysis of the b-factors of the three-dimensional structure of GA and molecular dynamics showed that this region had a high degree of lability. An increase in the thermal stability of a protein with mutation Thr390Ala had no good explanation in classical approaches, since the hydrogen bond between the hydroxyl group Thr390 and the carbonyl group Ser386 was lost. From the molecular dynamics data, we assume that this substitution reduced the local tension of the α-helix 12 (Fig. 4A), which led to stabilization of the C-terminus of the α-helix and preservation of its secondary structure at elevated temperatures (Fig. 4B). Also, a change in the secondary structure of this α-helix caused the stabilization of the loop involved in the catalytic center by forming two additional hydrogen bonds between: 1) the side chain of His391 and carbonyl group of Ala392; 2) the side chains of Tyr402 and Arg404. At the same time, the catalytic activity of GA variant Thr390Ala was higher than that of the wild type protein.

**Figure 4.**
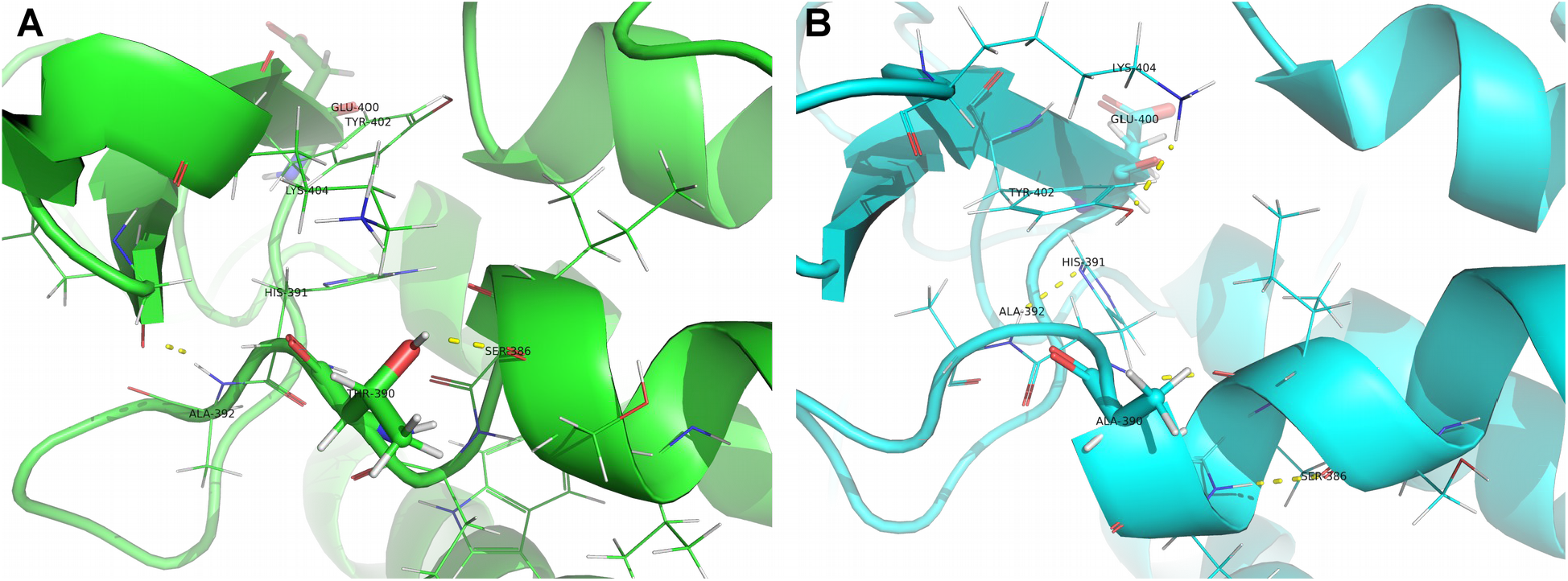
Amino acid substitution Thr390Ala. The structures of wild-type protein (A) and Thr390Ala mutant variant (B) are indicated. The Thr390Ala substitution reduced the local tension of the α-helix 12, which led to stabilization of the C-terminus of the α-helix and preservation of its secondary structure at elevated temperatures. Alteration in the secondary structure of α-helix caused the stabilization of the loop involved in the catalytic center by forming two additional hydrogen bonds between: 1) the side chain of His391 and carbonyl group of Ala392; 2) the side chains of Tyr402 and Arg404.

The mutant protein Thr390Ala/Ser436Pro had two amino acid substitutions. It was shown previously [29] that the substitution Ser436Pro led to thermal stabilization of GA. When comparing the data in Table 2 and the *k*_d_ dependency graphs from 1/T (Fig. 2B) for the mutant Thr390Ala and Thr390Ala/Ser436Pro variants, a weak additive effect of two mutations can be seen.

The GA double mutant Leu7Met/His391Tyr appeared to be much more stable than the wildtype protein. It is worth noting that the amino acid residues Leu7 and His391 do not interact with each other directly, however, they consist in the α-helices 1 and 12, respectively, which interact with each other through a set of amino acid residues: Leu3, Trp6, Leu7, Asp382, Val385, Ser386, Val388 and Glu389. Experiments have shown that a single mutant Leu7Met decreases the thermal stability of GA. According to the molecular dynamics data, this substitution upon thermal inactivation at 75°C leads to significant destabilization of the α-helices 1 and 13. It was shown that the His391Tyr substitution slightly increased the thermostability of GA, when another yeast, *S. cerevisiae*, was used as an expression system [13]. Results of molecular dynamics simulation confirms hypothesis about stabilization of glucoamylase by enhancement of hydrophobic interactions of aromatic tyrosine with amino acid residues Leu291, Ile387, Thr390, Phe402, and Gly407. It was shown that the change in ΔΔG^≠^ of this GA variant relative to the wild type protein was 1.3 kJ/mol at 65°C and only 0.2 kJ/mol at 80°C. The combination of the Leu7Met and His391Tyr mutations led to a significant thermal stabilization - the change in the free energy ΔΔG^≠^ at 80°C was 2.54 kJ/mol. This non-additive effect can be explained by a change in the protein denaturation pathway, where during the melting of the secondary structure of α-helix 1, the previously destabilizing Met7 fills the cavity between Val385, Leu422, Ala425 and Asn426 residues, which makes the molecule more compact and rigid, thereby stabilizing α-helices 1, 12 and 13 along with Tyr391.

The Ser8Arg/Gln338Leu double mutant also appeared to be more stable than the wild-type protein. When studying the thermostability of single mutations Ser8Arg and Gln338Leu, it turned out that the Gln338Leu substitution did not significantly reduce the thermostability of GA, and the Ser8Arg substitution was thermally stabilizing. The thermostability improvement of the Ser8Arg mutation can be explained by the stabilization of the α-helix 1 due to the formation of a salt bridge between the side chains of Arg8 and Asp4. However, the Ser8Arg mutant form of GA was less thermostable than double mutant Ser8Arg/Gln338Leu: the change in free energy ΔΔG^≠^ at 80°C was 1.06 kJ/mol for Ser8Arg, whereas for the Ser8Arg/Gln338Leu variant it reached value of 2.23 kJ/mol at the same temperature.

The similar situation was observed for the Asn9His/Ile82Phe mutant form. In this case, a Asn9His single substitution leads to destabilization of the GA due to the loss of the hydrogen bond between the side chain of Asn9 and the carbonyl group of Ser5. Substitution Ile82Phe increases the thermal stability of GA, which is achieved by enhancing the hydrophobic interactions of Phe82 in a pocket formed by amino acid side chains of Tyr81, Met135, Phe138, Ile154, Val155 and Ile158. However, as can be seen from the data in Table 2, the Ile82Phe variant is less thermostable than the Asn9His/Ile82Phe double mutant: ΔΔG^≠^ at 80°C is 1.08 kJ/ mol for Ile82Phe, while for the Asn9His/Ile82Phe variant - 2.0 kJ/mol at the same temperature. According to molecular dynamics data at 75°C for Ile82Phe and Asn9His/Ile82Phe variants, different thermal inactivation paths are observed. There is a loss of the secondary structure of the α-helices 2 and 3 from AAs 63 to 82 in the molten globule of the mutant variant Ile82Phe (Fig. 5). This molten globule state stabilized by hydrogen network between side chains of Ser34 and Glu79 and carbonyl and amine groups of Gly23, Ala24, Gly35 and Ile36 alpha chains. Meanwhile molten globule of Asn9His/Ile82Phe variant has different conformation where C-terminus of α-helix 2 and N-terminus α-helix 3 stabilized by newly formed hydrophobic core between side chains of Leu10, Leu19, Ile22, Val37, Val59, Leu60, Leu63, Leu74 amino acids and hydrogen bond network of Gly23, Ala24, Asp25, Asp33, Ser34, Gly35, Ile36, Asn80 and Glu113 side chains.

**Figure 5.**
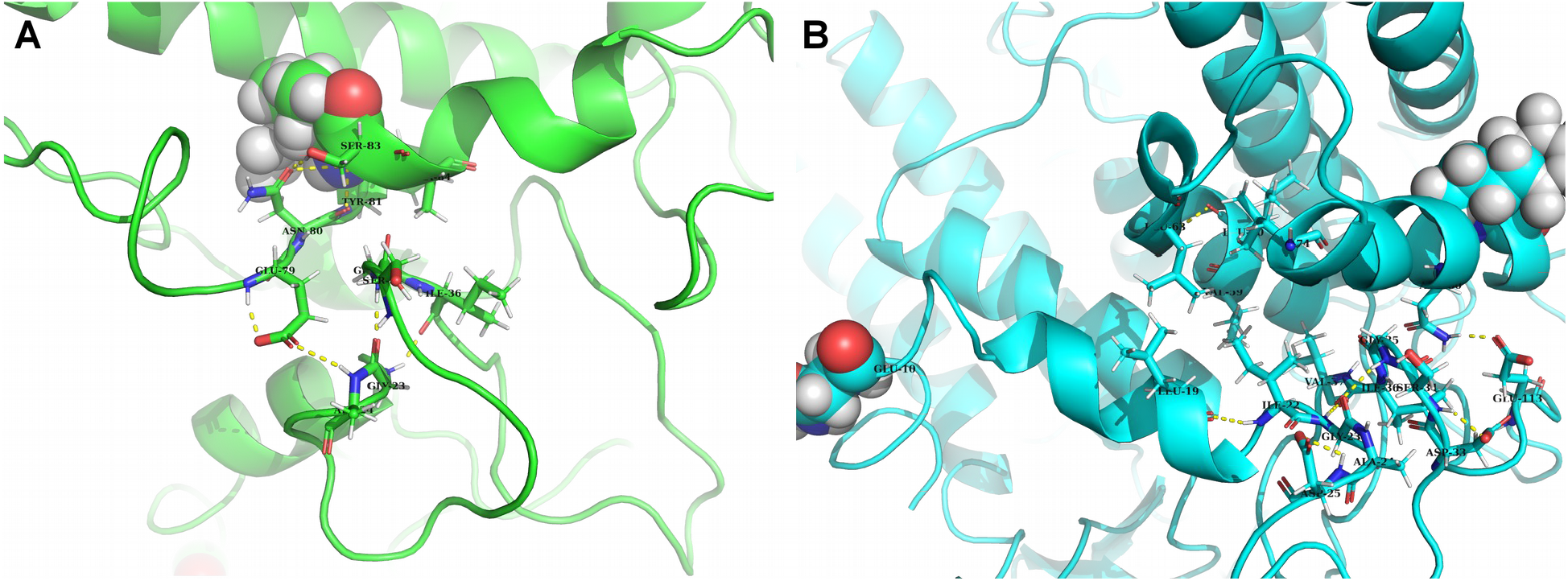
Amino acid substitutions Ile82Phe and Asn9His. The structures of Ile82Phe single mutant (A) and Asn9His/Ile82Phe double mutant (B) are indicated.

## Discussion

The methods of directed evolution can be used to obtain a protein with improved characteristics even if the structure and catalytic mechanism of enzyme are still unknown. However, this requires efficient methods for creating DNA libraries of mutant genes, an expression system that is suitable for the protein under study, and a reliable and fast method for clone selection. The disadvantage of a directed evolution strategy is the need for screening from thousands to tens of thousands of transformants.

In this work, the yeast *Pichia pastoris* GS115, which had a high level of glucoamylase production, was chosen as the expression system. This made it possible to immediately build up and analyze a large number of clones in small volumes of the culture medium. The pPEHα plasmid used here contained an alpha-factor that ensured the secretion of glucoamylase into the culture medium. The *HIS4* gene allowed selection of transformants on the basis of histidine independence, rather than zeocin resistance, which significantly reduced the cost of screening.

Unlike many other approaches, where all transformants were immediately tested for the presence of protein with the desired properties, we used a multi-stage screening. At the first stage, clones with glucoamylase activity were selected. Of the 3000 analyzed transformants, only 1000 were able to express active glucoamylase. The inability of others seemed to be as a result of the mutations leading to the protein was not expressed, either had an incorrect fold, or lost its catalytic activity.

A screening scheme for mutant gene libraries in 96- and 24-well plates was developed to select thermostable GA variants. The procedure had the advantage that after expression in the plate, the culture fluid could be used in several analyzes at once. Also, through the use of a plate photometer, we obtained quantitative, not qualitative, results when evaluating the thermal stability of clones. This increased the accuracy of the analysis and reduced the number of false-positive and false-negative tests.

In summary, the use of directed evolution approach in *Pichia pastoris* cells combining with the episomal expression vector pPEHα allowed us to obtain six new thermostable glucoamylase forms: Val301Asp, Thr390Ala, Thr390Ala/Ser436Pro, Leu7MetHis391Tyr, Asn9His/Ile82Phe, and Ser8Arg/Gln338Leu. A total of nine amino acid substitutions were found, seven of which, namely Val301Asp, Thr390Ala, Leu7Met, Asn9His, Ile82Phe, Ser8Arg, and Gln338Leu was new. As a result of the study of seven GA mutant forms with single amino acid substitutions, it turned out that only Val301Asp, Thr390Ala, Ile82Phe and Ser8Arg had increased thermal stability relative to wild type GA. In the case of the GA variants Leu7Met/His391Tyr, Asn9His/Ile82Phe, Ser8Arg/Gln338Leu, we observed the phenomenon of positive epistasis, when the combination of destabilizing and stabilizing substitutions led to a significant increase in the thermal stability of double mutant forms relative to the His391Tyr, Ile82Phe and Ser8Arg single mutants.

## Acknowledgements

Authors are very grateful to Andrey Komissarov for help with Sanger sequencing. The results of the work were obtained using computational resources of MCC NRC «Kurchatov Institute» and Peter the Great Saint-Petersburg Polytechnic University Supercomputing Center.

## Authors’ Contributions

## References

1. Reilly,PJ. (2010) In Whitaker,J.R., Voragen,A.G.J., Wong,D.W.S. (eds), Handbook of Enzymology. New York, Marcel Dekker, Inc., pp. 727–738.

2. Norouzian,D., Akbarzadeh,A., Scharer,J.M. and Young,M.M. (2006) Biotechnol. Adv., 24, 80–85.

3. Chen,H.-M., Li,Y., Panda,T., Buehler,F.U., Ford,C. and Reilly,P.J. (1996) Protein Eng., 9, 449–505.

4. Li,Y., Coutinho,P.M. and Ford,C. (1998) Protein Eng. 11, 661–667.

5. Liu,H.-L., Doleyres,Y., Coutinho,P.M., Ford,C. and Reilly,P.J. (2000) Protein Eng., 13, 655–659.

6. Allen,M.J., Coutinho,P.M., Ford,C. (1998) Protein Eng., 11, 783–788.

7. Surzhik,M.A., Churkina, S.V., Shmidt,A.E., Shvetsov,A.V., Kozhina,T.N., Firsov,D.L, Firsov,L.M. and Petukhov,M.G. (2010) Appl. Biochem. Microbiol,. 46, 221–227.

8. Surzhik,M.A., Schmidt,A.E., Glazunov,E.A., Firsov,D.L. and Petukhov,M.G. (2014) Appl. Biochem. Microbiol., 50, 118–124.

9. Liu,H.-L. and Wang,W.-C. (2003a) Biotechnol. Prog., 19, 1583–1590.

10. Liu,H.-L. and Wang,W.-C. (2003b) Protein Eng., 16, 19–25.

11. Aleshin,A.E., Golubev,A., Firsov,L.M. and Honzatko,R.B. (1992) J. Biol. Chem., 267, 19291–19298.

12. Aleshin,A.E., Firsov,L.M. and Honzatko,R.B. (1994) J. Biol. Chem., 269, 15631–15639.

13. Wang,Y., Fuchs,E., da Silva,R., McDaniel,A., Seibel,J. and Ford,C. (2006) Starch, 58, 501–508.

14. McDaniel,A., Fuchs,E., Liu,Y. and Ford, C. (2008) Microb. Biotechnol., 1, 523–531.

15. Sambrook,J. and Russel,D.W. (2001) Molecular Cloning: A Laboratory Manual, 3rd ed., Cold Spring Harbor Laboratory, Cold Spring Harbor, NY.

16. Wu,S. and Letchworth,G.J. (2004) Biotechniques, 36, 152–154.

17. Lin-Cereghino,J.L., Wong,W.W., Xiong,S., Giang,W., Luong,L.T., Vu,J., Johnson,S.D. and Lin-Cereghino,G.P. (2005) Biotechniques, 38, 44–48.

18. Innis,M.A., Holland,M.J., McCabe,P.C., Cole,G.E., Wittman,V.P., Tal,R., Watt,K.W.K., Gelfand,D.H., Holland,J.P. and Meade,J.H. (1985) Science., 228, 21–26.

19. Zheng,L., Baumann,U. and Reymond,J.L. (2004) Nucleic Acids Res., 32, e115.

20. Liu,H. and Naismith,J.H. (2008) BMC Biotechnol., 8, 91. doi:10.1186/1472-6750-8-91

21. Cadwell,R.C. and Joyce,G.F. (1994) Genome Res., 3, 136–140.

22. DeLano,W.L. (2014) The PyMOL Molecular Graphics System, Version 1.8. Schrödinger LLC.

23. Abraham,M.J., Murtola,T., Schulz,R., Páll,S., Smith,J.C., Hess,B. and Lindahl,E. (2015) SoftwareX, 1–2, 19–25.

24. Lindorff-Larsen,K., Piana,S., Palmo,K., Maragakis,P., Klepeis,J.L., Dror,R.O. and Shaw,D.E. (2010) Proteins, 78, 1950–1958.

25. Kirschner,K.N., Yongye,A.B., Tschampel,S.M., Gonzalez-Outeirino,J., Daniels,C.R., Foley,B.L. and Woods,R.J. (2008) J. Comput. Chem., 29, 622–655.

26. Goette,M. and Grubmüller,H. (2009 J. Comput. Chem., 30, 447–456.

27. Seeliger,D. and de Groot, B. L. (2010) Biophys. J., 98, 2309–2316.

28. Gapsys,V., Michielssens,S., Seeliger,D. and de Groot,B.L. (2015) J. Comput. Chem., 36, 348–354.

29. Li,Y., Reilly,P.J. and Ford,C. (1997) Protein Eng., 10, 1199–1204.

30. Urabe,I., Nanjo,H., and Okada,H. (1973) Biochim. Biophys. Acta, 302, 73–79.

